# Plant seeds are primed by herbivore-induced plant volatiles

**DOI:** 10.1101/522839

**Authors:** Abhinav K. Maurya, Leila Pazouki, Christopher J. Frost

## Abstract

Mature plants can detect and respond to herbivore-induced plant volatiles (HIPVs) by priming or directly activating defenses against future herbivores. Whether other plant life stages can respond to HIPVs in similar manners is poorly understood. For example, seeds are known to respond to a variety of environment cues that are essential for proper germination timing and survival. Seeds may also be exposed to HIPVs prior to germination, and such exposure may affect the growth, development, and defense profiles when the seeds grow into mature plants. Here, we investigated the effect of seed exposure to common HIPVs on growth, reproduction and defense characteristics in the model plants *Arabidopsis thaliana* and *Medicago truncatula*. Of all the HIPVs tested, indole specifically reduced both beet armyworm growth on *A. thaliana* and pea 33 aphid fecundity on *M. truncatula*. Induction of defense genes was not affected by seed exposure to indole in either plant species, suggesting that seed priming operates independently of induced resistance. Moreover, neither species showed any negative effect of seed exposure to HIPVs on vegetative and reproductive growth. Rather, *M. truncatula* plants derived from seeds exposed to *z*-3-hexanol and *z*-3-hexenyl acetate grew faster and produced larger leaves compared to controls. Our results indicate that seeds are sensitive to specific HIPVs, which represents a novel ecological mechanism of plant-to-plant communication.

## Introduction

Spermatophytes (or seed plants) are a dominant clade of vascular plants on earth (Friis *et al.*, 2011; Simonin & Roddy, 2018). Their dominance is due to large part to the evolution of the seed, which provides protection to the embryo prior to germination and nutrition during the transition to autotrophy. One advantage of the seed is the ability to survive long periods of time in dormancy until environmental conditions are suitable for germination and growth. During dormancy, seeds are inevitably exposed to a variety of biotic and abiotic environmental conditions such as temperature, moisture, fire, soil chemicals, and chemical exudates of plant and microbial origin that may affect their germination (Fenner, 2000). Many of these conditions are well-established cues that seeds use to coordinate their physiology and metabolism to properly time germination to maximize viability and establishment (Karssen & Hilhorst, 2000; Bentsink & Koornneef, 2008). Temperature (Probert, 2000; Reynolds *et al.*, 2001), rainfall (Gutterman, 1994; Pake & Venable, 1996; Levine *et al.*, 2008), and light (Wesson & Wareing, 1969; Milberg *et al.*, 2000; Flores *et al.*, 2006) are well-documented abiotic environmental cues that affect the germination of seeds, and responses to these cues are regulated through phytohormone signaling pathways (Chen *et al.*, 2008; Seo *et al.*, 2008; Toh *et al.*, 2008).

In addition to abiotic cues, seeds can perceive a variety of chemical cues of biological origins that can affect germination and subsequent defensive profiles. For example, low molecular weight phenolic compounds in soil (Muscolo *et al.*, 2001), artemisinin released from leaves (Chen & Leather, 1990) and catechin released from plants after herbivory (Thelen *et al.*, 2005) inhibit seed germination. In contrast, smoke-derived karrikins (Flematti *et al.*, 2004; Dixon *et al.*, 2009; Nelson *et al.*, 2012) and strigolactone (SL) phytohormones released from plant roots can stimulate seed germination (Cook *et al.*, 1966; Bergmann *et al.*, 1993). Moreover, recent studies have shown that seeds are receptive to the direct application of exogenous phytohormones that can activate plant defenses (Rajjou *et al.*, 2006; Worrall *et al.*, 2012; Jucelaine *et al.*, 2018). For example, treating tomato seed with the phytohormone jasmonic acid (JA) and β-aminobutryric acid (BABA) lead to JA- and ethylene-dependent resistance in future plants against spider mite, caterpillars, aphids, and pathogens (Worrall *et al.*, 2012). Seed treatment with JA also changes the volatile composition of mature plants, making their blends more attractive to predatory mites (Smart *et al.*, 2013). Similarly, seed treatment with salicylic acid (SA) enhances the expression of SA-related genes and the endogenous SA level against root holoparasite (*Orobanche cumana*) (Yang *et al.*, 2016). Additionally, seed coating with plant growth promoting rhizobacteria (PGPR) and plant growth promoting fungus (PGPF) enhances seed germination, seedling establishment, and boosts induced defenses in future plants in SA-, ET-, and JA-dependent manners (Ryu *et al.*, 2004; Rudrappa *et al.*, 2010; Sharifi & Ryu, 2016).

Seeds also come in contact with biotic agents that are volatile. Inhibitory and allelopathic effects of some plant and microbial derived volatile organic compounds (VOCs) have been known for a long time (Muller & Muller, 1964; Muller, 1965; Oleszek, 1987; Bradow & Connick, 1990; Koitabashi *et al.*, 1997; Mirabella *et al.*, 2008). Whereas these VOCs do not necessarily provide contextual information about future environmental conditions, herbivore-induced plant volatiles (HIPVs) represent potentially reliable and adaptive indicators of herbivory. The function of HIPVs in priming or directly inducing plant defenses is now well established (Engelberth *et al.*, 2004; Frost *et al.*, 2007; Rodriguez-Saona & Frost, 2010), and exposure of undamaged plants to HIPVs is known to induce or prime the genes in phytohormone pathways (Bate & Rothstein, 1998; Engelberth *et al.*, 2007; Frost *et al.*, 2008c). Moreover, aboveground HIPV priming cues are also produced belowground by plant roots (Lawo *et al.*, 2011; Palma *et al.*, 2012; Gfeller *et al.*, 2013; Barsics *et al.*, 2017) and rhizosphere organisms (Bhattacharyya *et al.*, 2015; Kanchiswamy *et al.*, 2015). Therefore, there are multiple routes by which seeds could be exposed to HIPVs, including simple diffusion of HIPVs produced belowground (Peñuelas *et al.*, 2014) and precipitation and leaching of HIPVs produced aboveground (Muller *et al.*, 1964; H B Tukey, 1970). While some HIPVs may have allelopathic effects on seed germination (Preston *et al.*, 2002; Karban, 2007; Mirabella *et al.*, 2008), whether exposure of seeds to HIPVs alters subsequent plant physiology and defense is currently unknown.

Here, we determined the effect of seed exposure to HIPVs on plant growth and direct defenses. Specifically, we used a comparative approach to investigate the effects of HIPV exposure to the seeds of (1) *A. thaliana* on the performance of a chewing herbivore (beet armyworm; *Spodoptera exigua*) and (2) *M. truncatula* on the performance of a phloem feeding herbivore (pea aphid*; Acyrthosiphon pisum*). We also tested the effect of seed exposure to plant volatile on the growth, development, and defense gene expression of *A. thaliana* and *M truncatula*. We specifically tested HIPVs that have been shown previously to prime mature plants: indole, *cis*-3-hexenol (*z*3HOL), *cis*-3-hexenyl acetate (*z*3HAC), β-caryophyllene (BCP), and *trans*-2-hexanol (*e*2HAL). We predicted that HIPV exposure to seeds would prime the resulting mature plants for enhanced resistance against both chewing and phloem-feeding herbivores.

## Materials and Methods

### Plant material

*A. thaliana* (Col-0) seeds were surface sterilized in 75% (v/v) ethanol for five minutes and 20% bleach (v/v) in .01% Tween-20 for ten minutes. After sterilization the seeds were washed three times with distilled water and spread on petri-plates with wet Whatman paper. Petri plates were kept at 4°C for 2 days, this allowed the seeds to break dormancy and synchronize germination.

*M. truncatula*, A-17 seeds were scarified in concentrated H_2_SO_4_ for 10 min and surface sterilized in 20% (v/v) bleach in 0.1% (v/v) Tween-20 solution for 10 min. Seeds were rinsed five times with sterile water and were spread on petri plates with wet Whatman paper. Petri plates were covered with aluminum foil to maintain dark environment and kept at 4°C for two days.

### Seed treatment with plant volatiles

Volatile dispensers were used to treat *A. thaliana* and *M. truncatula* seeds to individual plant volatiles. For preparing volatile dispensers 20 μl of *cis*-3-hexenol, *cis*-3-hexenyl acetate (Engelberth *et al.*, 2004), trans-2-hexenal, β-caryophyllene and 20 mg indole (Erb *et al.*, 2015) was added into separate 2.0 ml amber glass vial (Agilent Technologies) with 1 mg of glass wool (Fig. S1). Control volatile dispensers had only glass wool without any volatile. The amber vials with volatiles were sealed with a rubber septum and connected to the 2-ounce plastic cup by piercing the plastic cup and amber vial rubber septum with an 18-gauge needle. This procedure is similar to what has been used previously for controlled administration of HIPVs (Erb *et al.*, 2015). Each volatile was administered to seeds in multiple plastic cups (biological replicates) and number of seeds planted from each plastic cups constituted the technical replicates.

### *A. thaliana* Seed germination

Each volatile was administered to seeds in 5 replicates (plastic cups). After one day of volatile treatment, two *A. thaliana* seeds were transferred from each plastic cups to agar plates containing 1.0% (w/v) agar (Sigma) and standard 0.5X MS medium (Murashige and Skoog basal at an adjusted pH of 7.0). Total 9 agar plates were used for each volatile treatment. The Petri dishes were kept in growth chamber at 25°C under a 16 h light: 8 h dark (16L: 8D) day/night cycle for two days. Percent seed germination was measured after two days of seed transfer from plastic cup to petri-plates.

### *A. thaliana* growth

After one day of volatile treatment, *A. thaliana* seeds were transferred to 5.5 x 5.5 x 5.5 cm pots filled with sterile Metro-Mix 360 soil. After transplanting, pots were placed on trays (54 x 28 x 6 cm).in a growth chamber at 25°C under a 12 h light: 12 h dark (12L: 12D) cycle. Once germinated seedlings reached to 4-6 leaf stage, they were fertilized twice a week with 10 ml 1/2 strength Hoagland’s solution. Arabidopsis growth and fitness were measured in terms of number of leaves, maximum rosette diameter, the length of the bolt and number of siliques produced.

### *M. truncatula* growth

Volatile exposed *M. truncatula* seeds were planted in 9 x 6.5 x 6.5 cm pots as described above. The trays were kept in growth chamber at 25°C under a 12 h light: 12 h dark (12L: 12D) day/night cycle for ten days. After 10 days the trays were moved to green house and kept there till the end of the experiment. *M. truncatula* growth and fitness were measured in terms of petiole length, leaf blade length, leaf blade width, main shoot length, axillary shoot length and number of fruits using numerical nomenclature coding system developed by Bucciarelli *et al.* (2006). The numerical nomenclature for vegetative growth (Fig. S3) starts with first unifoliate leaf as metamer 1 (m1) followed by first trifoliate as metamer 2 (m2) and so on. The axillary shoots are coded as per the their metamer of origin (e.g. the axillary shoot originating from first unifoliate or metamer 1 is also designated as m1). Additionally, decimal addition to numerical coding system defines the development stage of leaf (e.g. m2.1 represent the bud break for the first trifoliate, m2.5 represent the half open blade of first trifoliate while m2.9 represent fully developed first trifoliate).

### Caterpillar herbivory

Beet armyworm (*Spodoptera exigua*) was used to evaluate the effect of seed exposure to HIPVs on herbivore defense of Arabidopsis plants. Caterpillar eggs were ordered from Benzon Research Inc. USA (Permit #P526P-16-02563). Egg masses were immediately transferred to artificial diet in 2-ounce plastic cups. Eggs in plastic cups were maintained at 24°C on artificial diet until the desired instar. Third instar caterpillars were used for feeding experiment on five to six-week-old, vegetative stage, *A. thaliana* plants. For the first feeding experiment, each volatile was administered to seeds in six plastic cups (biological replicates) and three seeds were planted from each plastic cups (three technical replicates). For the second feeding experiment, each volatile treatment had 10 biological replicates) and three technical replicates. For feeding experiment caterpillars were starved for 3 hours and weighed before their transfer to Arabidopsis plants. One third-instar caterpillar was placed on each Arabidopsis plants. The plants were covered with a nylon mesh bag to avoid the caterpillar escape. The caterpillars were allowed to feed freely for 24 h before being removed from the plants. After their removal, the caterpillars were kept at room temperature for three hours to allow the digestion of ingested plant material. Caterpillars that molted during the second experiments were removed from the assay analysis. After 3 h the caterpillars were weighted on microbalance. Aboveground plant material was also collected in liquid nitrogen and stored at -80 °C for later molecular work.

### Aphid herbivory

Pea aphid (*Acyrthosiphon pisum*) colony was maintained on fava bean plant kept in growth chamber (20 °C, 12:12 h light:dark). For aphid feeding experiment, three adult aphids (defined as F_0_ generation) (Tomczak & Müller, 2017) were placed in an insect bag (L15 X W6, BugDorm) on three trifoliate (8 to 10 plants per treatment). After 24 h, the adults were removed and one trifoliate was collected while 5 nymphs (defined as F_1_ generation) were left on the plant for 13 more days. For 13 days the nymphs grew and produced offspring (F_2_ generation). On 14^th^ day the all the aphids were collected, the total offspring (F_2_) were counted and weighed on microbalance. Aboveground plant material was also collected on day 14 in liquid nitrogen and stored at -80 °C for later molecular work.

### Gene expression analysis

Aboveground tissue collected from *A. thaliana* plants after one day of caterpillar herbivory and *M. truncatula* after 14 days of aphid feeding were used for gene expression analysis. Total RNA was isolated from approx. 150 mg of ground tissue using modified cetyl trimethylammonium bromide (CTAB) method (Frost *et al.*, 2012). RNA was quantified with Nanodrop and integrity was confirmed using a native 1% agarose-0.5x TAE gel. Total RNA (2.5 μg per sample) was treated with DNAse (Turbo DNAse, Ambion), then 0.7 μg of DNA-free RNA was reverse-transcribed to cDNA using High Capacity cDNA Reverse Transcript Kit (Applied Biosystems). Real‐time PCR was done using the Quant Studio-3 PCR System (Applied Biosystems) with each reaction containing 2 μl of EvaGreen^®^ PCR Master Mix (Mango Biotechnology), 0.3 μl of 10 μM forward and reverse primer, 5.4 μl of DI water, and 2μl (2.5 ng) of cDNA in a total volume of 10 μl. Primer specificity was confirmed by melting curve analysis, and relative transcript levels were calculated using the 2^-∆CT^ method (Livak & Schmittgen, 2001) with elongation factor 1-alpha (*EF1-α*) and Glyceraldehyde 3-phosphate dehydrogenase (*GAPDH*) as reference genes for *M. truncatula* and Actin-7 and *GAPDH* as reference genes for *A. thaliana*. Primer sequences for all *M. truncatula* and *A. thaliana* genes tested are listed in Table 1.

**Table 1.**
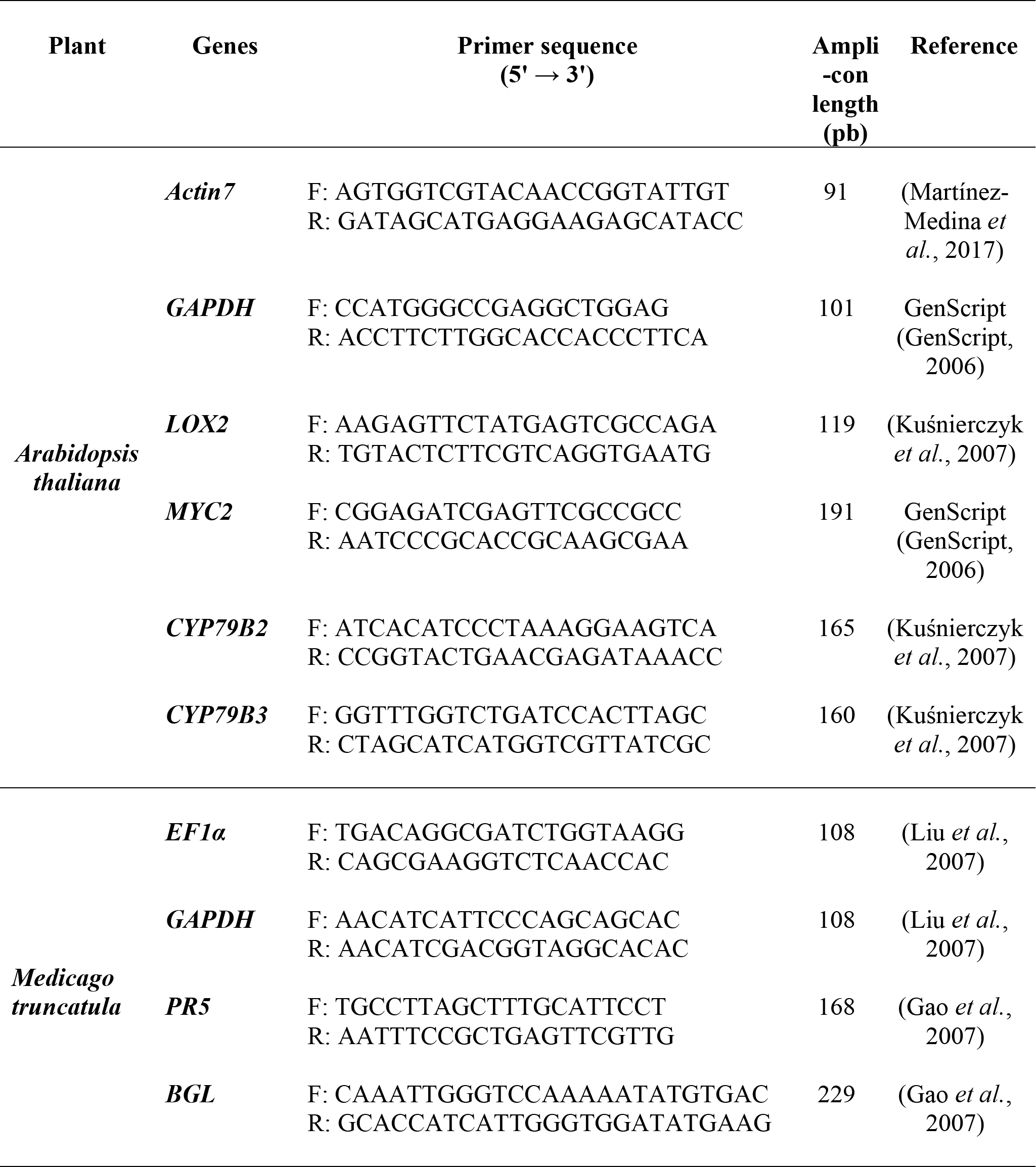
Primer sequences used in this study.

### Statistical analyses

Raw data were checked for normality and homogeneity of variance before performing the parametric tests. For *A. thaliana,* leaf number, rosette diameter, were analyzed using repeated measures ANOVA. For *M. truncatula*, leaf petiole length, leaf blade length and width, main shoot and axillary shoot length were analyzed using one-way ANOVA followed by Dunnett post-hoc test. Other response variables for *A. thaliana* and *M. truncatula* growth along with caterpillar growth rate, aphid number and aphid weight were analyzed for significance using student’s t-test. For t-test, treatments were compared to controls. The gene expression data were analyzed using one way ANOVA followed by Tukey’s post-hoc test. Statistical analyses were performed using R version 3.4.2 and GraphPad Prism and figures were generated via GraphPad Prism.

## Results

### Seed exposure to indole enhances plant resistance against chewing and sap feeding herbivores

Indole exposure to seeds reduced the relative growth rate of *S. exigua* caterpillars feeding on mature foliage by 33% (p=0.0706, Fig. 1a) and 30% respectively (p=0.0124, Fig. 1b) in separate experiments. In contrast, seed exposure to GLVs and terpenes had no effect on caterpillar growth (p>0.05, Fig. 1a). We observed similar effects of indole in *M. trucatula*, where pea aphids fecundity and total weight were reduced by 28% (p=0.007, Fig. 1c) and 41% (p=0.015, Fig. 1d), respectively. Additionally, *z*3HAC seed treatment to *M.trucatula* reduced pea aphid fecundityby 27% (p=0.0354 Fig. 1c) and total nymph weight by 35% (p=0.067 Fig. 1d).

**Fig. 1.**
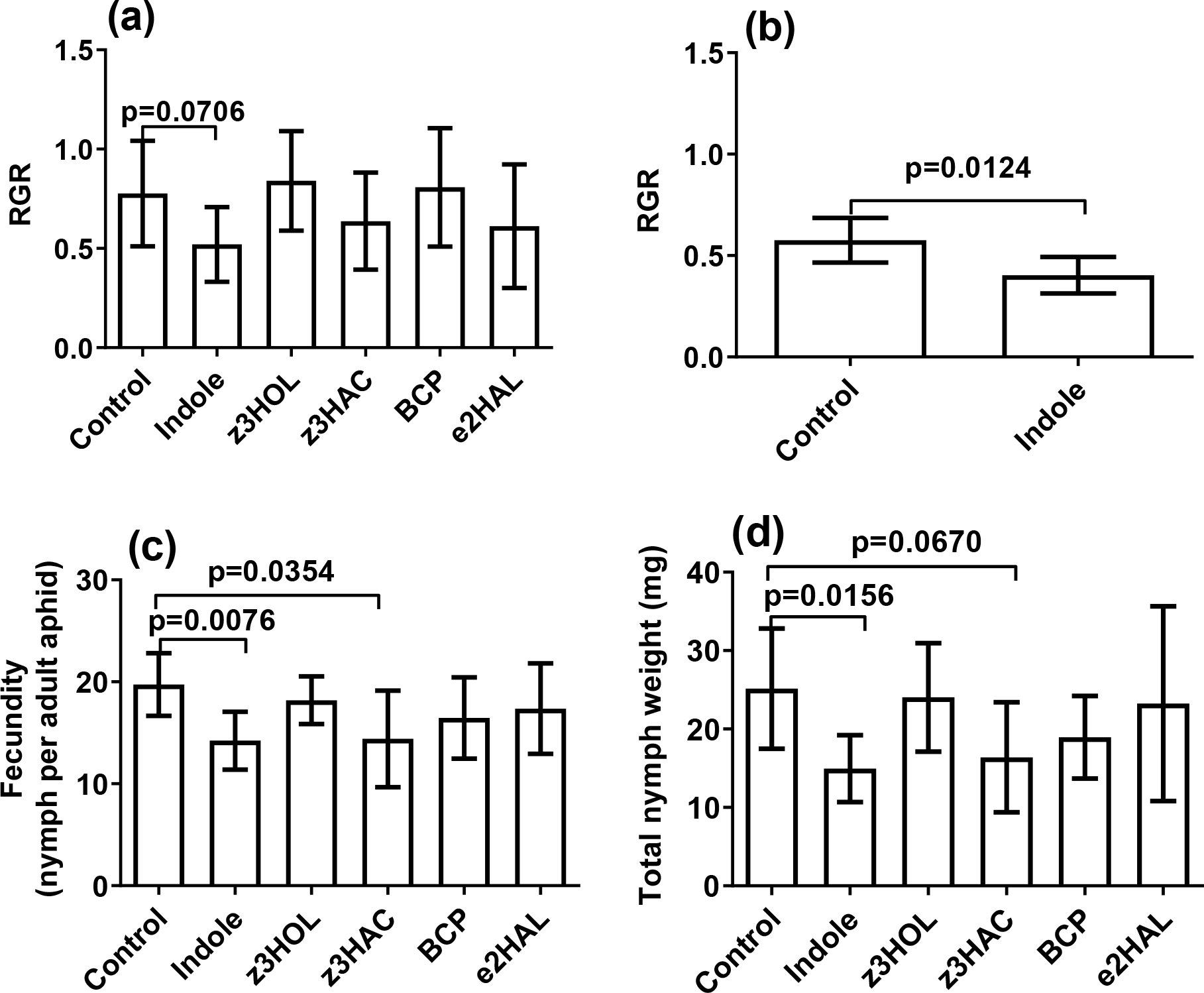
The effect of seed exposure to plant volatiles on the herbivore fitness (a) Relative growth rate (RGR) of *S. exigua* caterpillars after 24 h herbivory on *A. thaliana* plants grown from control and volatile-exposed seeds (*n*= 6, each biological replicate had 1-3 technical replicates), (b) Relative growth rate of *S. exigua* caterpillars after 24 h herbivory on *A. thaliana* plants grown from control and indole-exposed seeds in a separate caterpillar herbivory experiment (*n*=8-10, each biological replicate had 1-3 technical replicates), (c) Fecundity (nymph number per adult) and, (d) nymph weight after 14 days of herbivory on *M. truncatula* plant grown from control and volatile-exposed seed (n=6-8). Values are shown as means ± 95% CI and significance was calculated by student’s *t*‐test (two‐tailed).

### Seed exposure to indole does not affect growth and development of *A. thaliana*

*A. thaliana* seed exposure to HIPVs had no significant differences relative to controls on the vegetative and reproductive growth. We found no differences in leaf number (p_trt_= 0.997, Fig. 2a), rosette diameter (p_trt_=0.672, Fig. 2b), bolt length (p=0.333, Fig. 2c), silique number (p=0.460, Fig. 2d), and fresh shoot weight (p=0.107, Fig. 2e) of plant that were grown from seeds exposed to any HIPV relative to control plants.

**Fig. 2.**
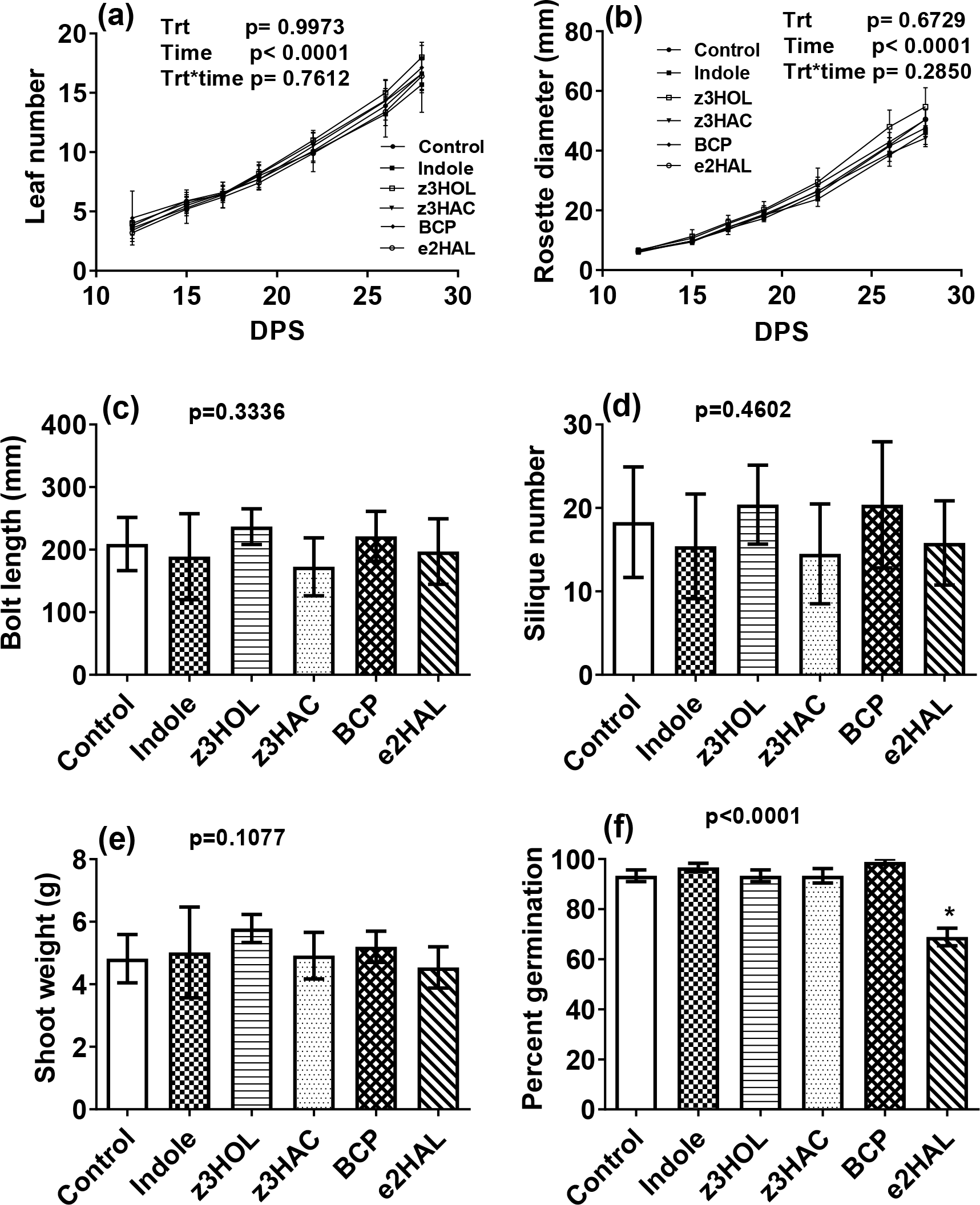
Seeds exposure to plant volatiles does not affect *A. thaliana* plants growth and reproductive output. The effect of seed exposure to plant derived volatiles on (a) Leaf number, (b) Rosette diameter, (c) Bolt length, (d) Mean silique number and (e) Shoot weight of plants. DPS represents days after seed sowing. Values are shown as means ± 95% CI (*n* = 8-10). Seed exposure to *e*2HAL reduced the seed germination on agar plates (e) Percent seed germination. Values are shown as means ± SEM (n=90). Significance was calculated by repeated measures ANOVA and one-way ANOVA.

We also measured the effect of HIPV exposure on seed germination of *A. thaliana* on MS media. Of all the HIPVs tested, only seed exposure to the GLV *e*2HAL reduced seed germination by 26% compared to control seeds (p<0.001, Fig. 2f).

### Seed exposure to GLVs enhances*M. truncatula* growth

*M. truncatula* seed exposure to *z*3HOL and *z*3HAC increased plant vegetative growth (Fig. 3a). Petiole length (p<0.05, Fig. 3b), leaf blade length (p<0.05, Fig. 3c), leaf blade width (Fig. 3d) and axillary shoot length (p<0.05, Fig. 3e) of the *z*3HOL and *z*3HAC exposed seed plants were higher compared to control plants while no such effect was seen on main shoot length (p_global_=0.016, p_Dunnett’s_>0.05, Fig. S2a). No other HIPV affected vegetative growth in *M. truncatula*. Furthermore, while *z*3HOL and *z*3HAC affected the vegetative growth, there was no difference in reproductive output of plants grown from HIPV-exposed seeds than control seeds (p=0.929, Fig. S2b).

**Fig. 3.**
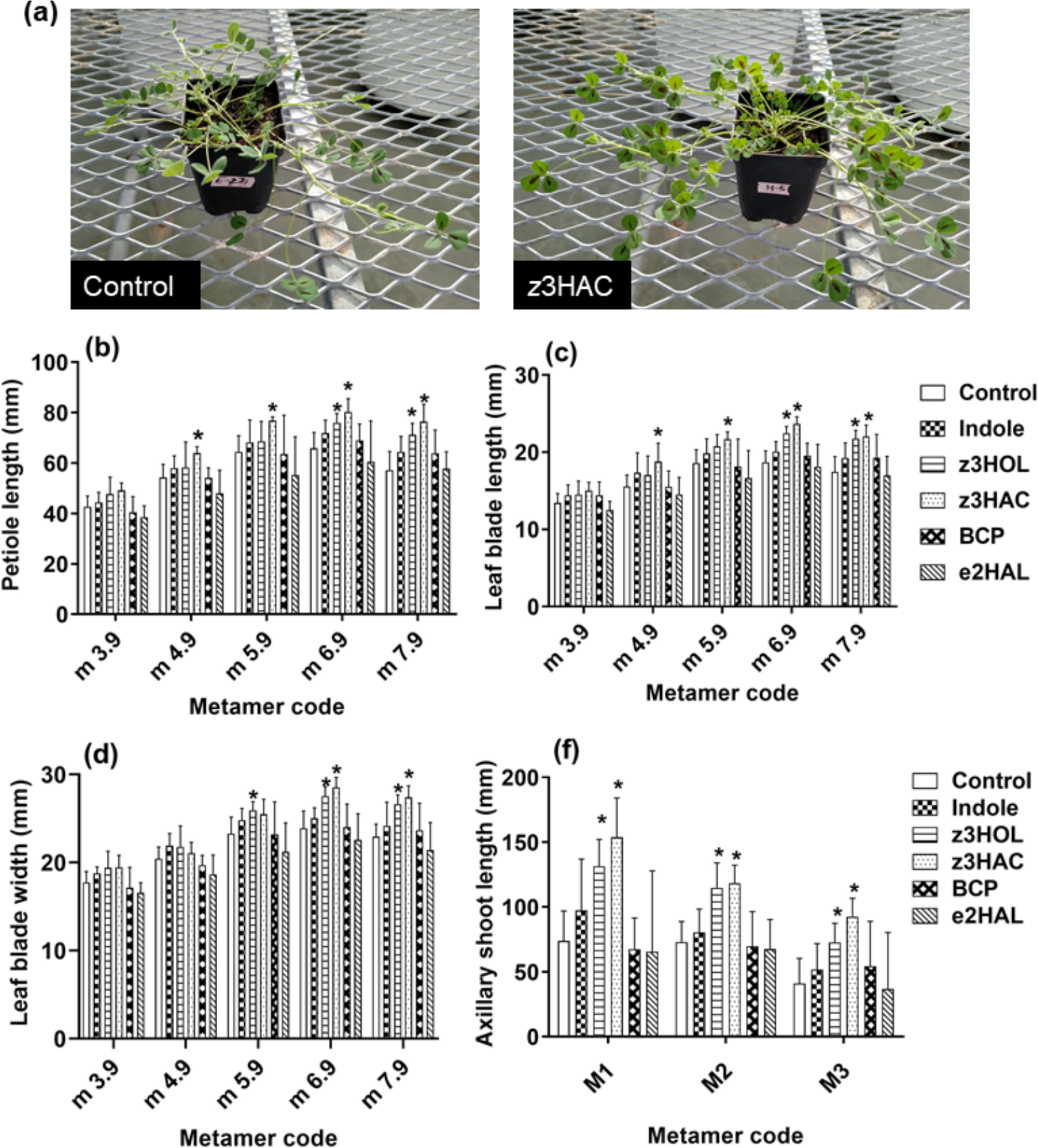
Seeds exposure to *cis* configuration green leaf volatiles enhances growth of *Medicago truncatula*. (a) Picture of control and *z*3HAC seed exposed *M. truncatula* plants. The effect of seed exposure to plant derived volatiles on (b) leaf petiole length, (c) Leaf blade length, (d) Leaf blade width and, (e) axillary shoot length. For leaf petiole length, leaf blade length and width all the measurements were taken when the leaves were fully developed. Axillary shoot was measured at 64 days after seed sowing. Values for each metamer are shown as means + 95% CI (*n*=5-10) and asterisks represent significant differences (p<0.05) from controls based on one-way ANOVA followed by Dunnett’s post-hoc analysis

### Seed exposure to indole does not affect herbivore-inducible defense gene expression after caterpillar or aphid herbivory

Since there was a clear effect of indole seed treatment on caterpillar and aphid fecundity, we assessed whether this effect was due to indole-mediated changes in inducible defenses. In *A. thaliana* challenged with *S. exigua*, we analyzed the expression of genes related to JA synthesis (*LOX2*, Fig. 4a) and signaling (*MYC2*, Fig. 4b), and glucosinolate biosynthesis (*CYB79-B2* and *CYB79-B3*, Fig. 4c-d). Caterpillar herbivory induced the expression of these four marker genes as expected, but indole-seed treatment neither directly stimulated nor statistically altered the caterpillar-induced expression patterns of these genes. In *M. truncatula* challenged with aphids, we analyzed two SA-regulated marker genes, *PR5* and *BGL-1*, which have previously been shown to be responsive to aphid feeding (Moran & Thompson, 2001; Gao *et al.*, 2008). *PR5* and *BGL-1* were induced by aphid feeding (Fig. 4e-f), but indole seed treatment neither directly stimulated nor statistically altered the aphid-induced expression patterns of these genes. That is, in all cases, indole did not directly induce, indirectly prime, or affect the magnitude of herbivore induction of these defense genes.

**Fig. 4.**
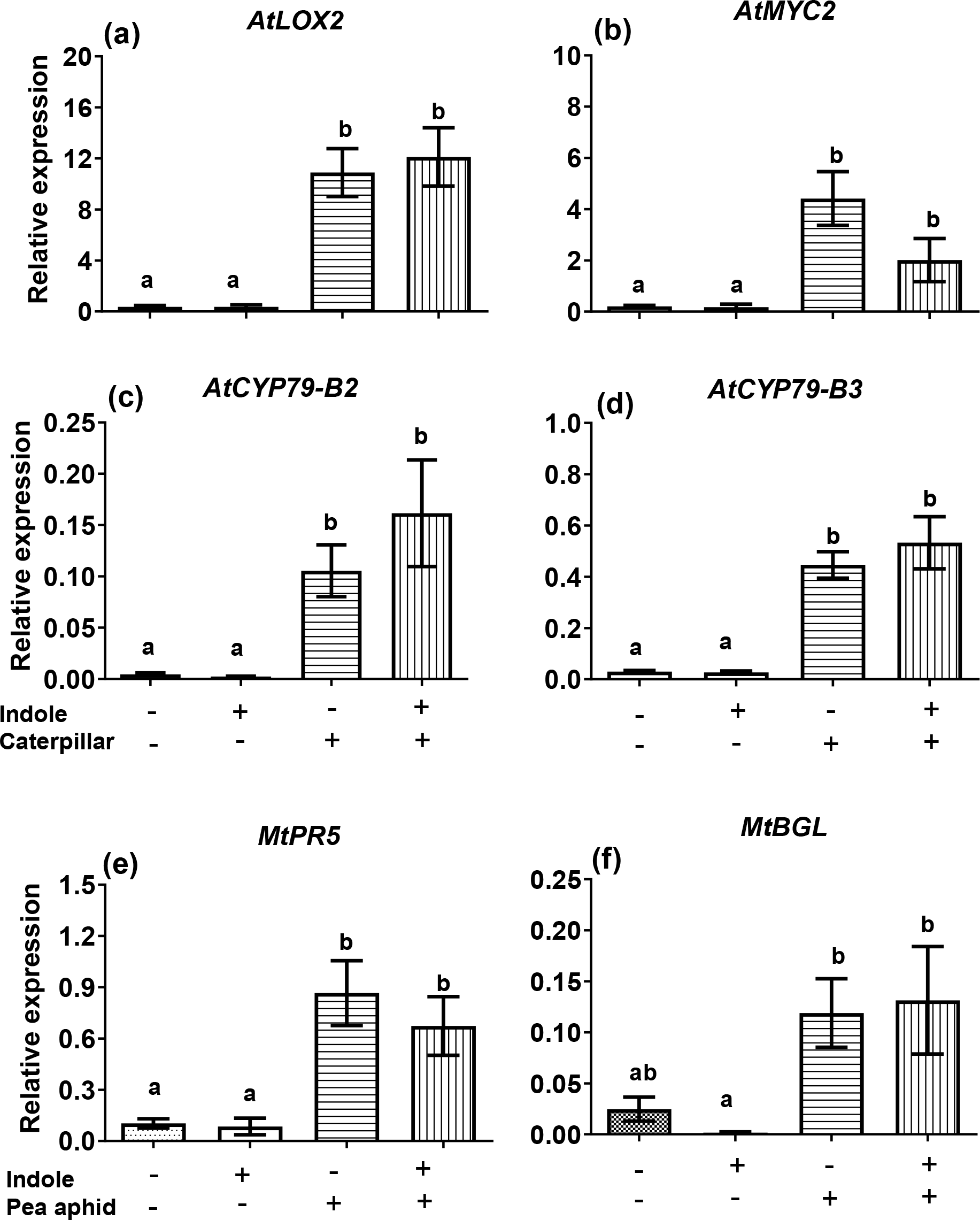
Seed treatment with indole does not enhance herbivore-induced expression of defense marker genes. Relative transcript levels of the genes *LOX2*, *MYC2*, *CYB-B2* and *CYB-B3* in *A. thaliana* after 24 h of *S. exigua* herbivory was measured by quantitative RT-PCR analysis (a-d). Similarly, transcript levels of SA regulated marker genes *PR5* and *BGL* were measured in *M. trunacatula* after 14 days of pea aphid herbivory (e & f). Relative expression was determined (2^-ΔCt^) using the geometric mean of two housekeeping genes for normalization. Bars represent mean ± SEM determined from three-five biological replicate assays, each biological replicate had two technical replicates. Different letters on the bar represent significant difference (p<0.05).

## Discussion

We show that seeds are viable receivers of HIPVs in ways that prime defenses and, in some cases, directly stimulate growth. Specifically, our study demonstrates that the pre-germination exposure of seeds to indole enhances resistance against herbivores of two feeding guilds in two different plant species without any apparent effects on plant growth or fitness. Our study also showed that seed exposure to *z*3HOL and *z*3HAC can enhance plant growth in *M. truncatula*. Biotic cues that reliably indicate future biotic stress can prime plant defenses for faster and/or stronger defenses following subsequent stress events (Conrath *et al.*, 2006; Frost *et al.*, 2008a). The phenomenon of HIPV-mediated priming is now well established in mature plants (Engelberth *et al.*, 2004; Frost *et al.*, 2007; Frost *et al.*, 2008b; Frost *et al.*, 2008c; Rodriguez-Saona *et al.*, 2009; Erb *et al.*, 2015). To our knowledge, our study is the first to show that seeds can also be primed by HIPVs. Moreover, seed exposure to HIPVs had no adverse effect on seed germination, vegetative growth and reproductive output of the primed mature plants (Fig. 2 & 3). Such a long-persisting defense response without apparent negative consequence on plant growth and development may be indicative of defense priming rather than direct activation of induced defenses.

HIPV-mediated defense priming is theoretically a component of an inducible resistance phenotype (Frost *et al.*, 2008a; Hilker *et al.*, 2016). Since seed treatment with defense phytohormones (e.g., JA, SA and BABA) primes defenses by modulating stress-related signaling pathways (Azooz, 2009; Worrall *et al.*, 2012; Jucelaine *et al.*, 2018), we hypothesized that volatile indole would prime seeds through inducible signaling pathways. We therefore predicted that seed-primed plants would show primed inducible defenses compared to controls when challenged with herbivores. For example, Worrall *et al.* (2012) showed that seed treatment with JA and BABA primed the antiherbivore and antipathogen defenses in mature Arabidopsis plants by JA-dependent processes. However, in our case, JA-related octadecanoid pathway (Wasternack, 2007; Ballaré, 2011) and glucosinolate biosynthesis (Hopkins *et al.*, 1998; Reymond *et al.*, 2004) marker genes were induced by *S. exigua* feeding to similar levels independent of indole seed treatment (Fig. 4). Similarly, marker genes for SA-related defense (Walling, 2008) in *M. truncatula* were induced by *A. pisum* but were not additionally enhanced by seed treatment (Fig. 4). Therefore, HIPV-mediated seed priming operates through a mechanism independent of inducible resistance. Moreover, indole seed treatment did not directly induce any marker gene before herbivory, further ruling out direct activation of induced resistance via seed priming (Fig. 4). Since we measured just single time points as indicators of inducible defense, it is possible that seed priming altered the temporal dynamics of induced defense. However, the time points we chose are reflective of sustained defense activation, which is one important aspect of defense priming. The enhanced defense in indole-exposed seed plants in our study is therefore likely a result of change in plant nutritive and defense chemistry.

Indole was the only HIPV we tested that primed plant defenses after seed exposure, and this effect was consistent across two model plants against herbivores of different feeding guilds. Indole is an ubiquitous, inter-kingdom intermediate in critical biochemical pathways (Zhang *et al.*, 2008) and a signaling molecule (Lee *et al.*, 2015). In plants, indole is also a common HIPV that contributes to direct and indirect defenses (Veyrat *et al.*, 2016; Gasmi *et al.*, 2018) and also acts as a defense priming cue (Erb *et al.*, 2015; Ye *et al.*, 2018). Our study adds an additional facet to the ecological role of indole in plant communication. That said, rhizosphere inhabiting bacteria also produce volatile indole, which can modulate plant growth via auxin pathway (Blom *et al.*, 2011; Yu & Lee, 2013; Bailly *et al.*, 2014; Bhattacharyya *et al.*, 2015). We tested the genes *CYP79B2* and *CYP79B3* in *A. thaliana* which involve in enzyme production that convert tryptophan (Trp) to indole-3-acetaldoxime (IAOx), a rate determining intermediate in auxin biosynthesis pathway and plant defense compound indole glucosinolates biosynthesis (Zhao *et al.*, 2002). Seed exposure to indole alone did not upregulate either gene, but *S. exigua* feeding induced their expression independent of seed exposure to indole (Fig. 4. c & d). Therefore, the auxin pathway may not be involved in indole-mediated seed priming. Nevertheless, seed priming was consistent in two different plant species against different feeding guilds of herbivores, suggesting a clear role for indole in mediating plant-seed communication.

Exposure of *M. truncatula* seeds to two GLVs (*z*3HOL and *z*3HAC) stimulated vegetative growth. Our group has recently seen similar vegetative and reproductive growth stimulation using a low-dose, persistent application of *z*3HAC in lima bean plants (Freundlich & Frost, 2018). In lima bean and *M. truncatula*, plants with increased growth also were better defended ((Freundlich & Frost, 2018) and Figs 1&3). GLVs are well-established priming cues against biotic stress (Engelberth *et al.*, 2004; Frost *et al.*, 2008c), and volatile communication between plants can alter biomass allocation (Ninkovic, 2003). Our results suggest that GLVs can also stimulate plant growth and ostensibly overcome the growth-defense dilemma (Herms & Mattson, 1992) in some plant species. One caveat, though, is that our group also has shown that persistent exposure to *z*3HAC reduces growth in *Capsicum annuum* (Freundlich & Frost, 2018), therefore the stimulating effect of GLVs is not universal.

As a final point, our results have potential applications in pest control and seed management. Recent attention has focused on leveraging priming of innate plant immunity (Pichersky & Gershenzon, 2002; Dervinis *et al.*, 2010; Song & Ryu, 2013; Song *et al.*, 2015; Pickett & Khan, 2016), due in part to presumed lower fitness costs of priming based defenses (van Hulten *et al.*, 2006; Buswell *et al.*, 2018). In-field foliar or soil application of these agents can induce plant defenses against herbivores (Bruce *et al.*, 2003; War *et al.*, 2011; Song & Ryu, 2013), but can also be prohibitively costly for large-scale application. In contrast, seed treatments are a common method of inoculating crops (Paparella *et al.*, 2015), and direct application of HIPVs to seeds could provide more viable priming-mediated solution to pest management. Moreover, *M. thaliana* is a close relative of fodder crop alfalfa, and improved vegetative growth after seed treatment with GLVs may provide a mechanism for enhancing fodder capacity and rejuvenating soils during crop rotations. Furthermore, HIPV-mediated seed priming may be a valuable tool in conservation efforts for rare or endangered species (Laetz *et al.*, 2009),if HIPV-mediated seed priming can enhance their innate immunity. Ultimately, seed priming via HIPVs represents a novel mechanism in plant-plant communication that may have trans-generational effects on ecological communities.

## Acknowledgements

This work was supported financially by a National Science Foundation grant IOS-1656625 to C.J.F. We thank Dr. Julia Frugoli for generously providing M. truncatula, A-17 seeds and Dr. Susana Karen Gomez for supplying the pea aphids. We are thankful to Grace Freundlich, Allie Peot and Rachel Haslem for lab assistance.

## Author Contributions

C.J.F. and A.K.M. designed the research. A.K.M. conducted experiments. A.K.M. and L.P. performed quantitative RT-PCR. A.K.M. and C.J.F. analyzed data and wrote the manuscript. All authors read and approved the manuscript.

## Supplemental Information

**Fig. S1.**
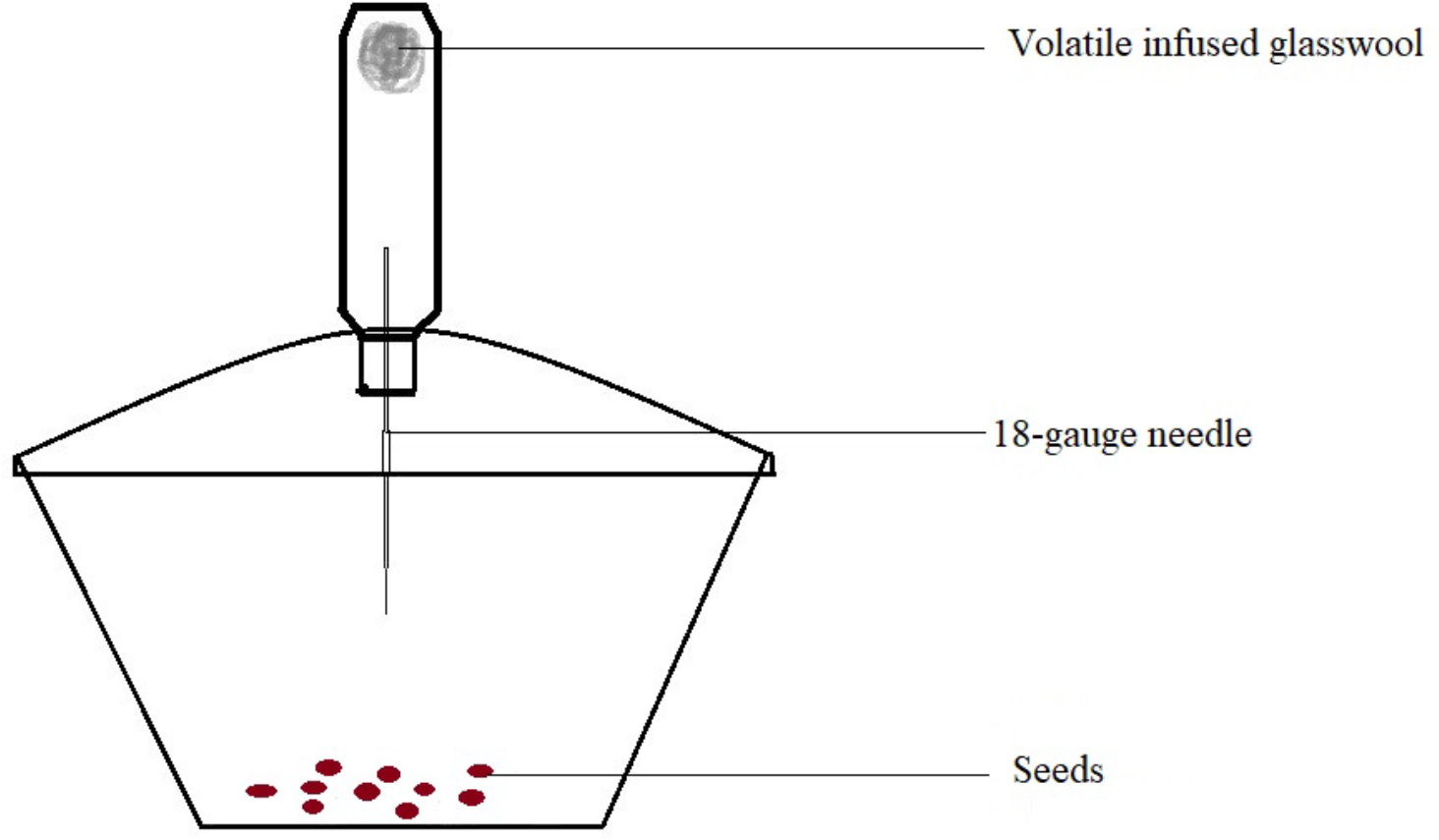
Pictorial representation of volatile dispensers used to expose seeds to synthetic plant volatiles.

**Fig. S2.**
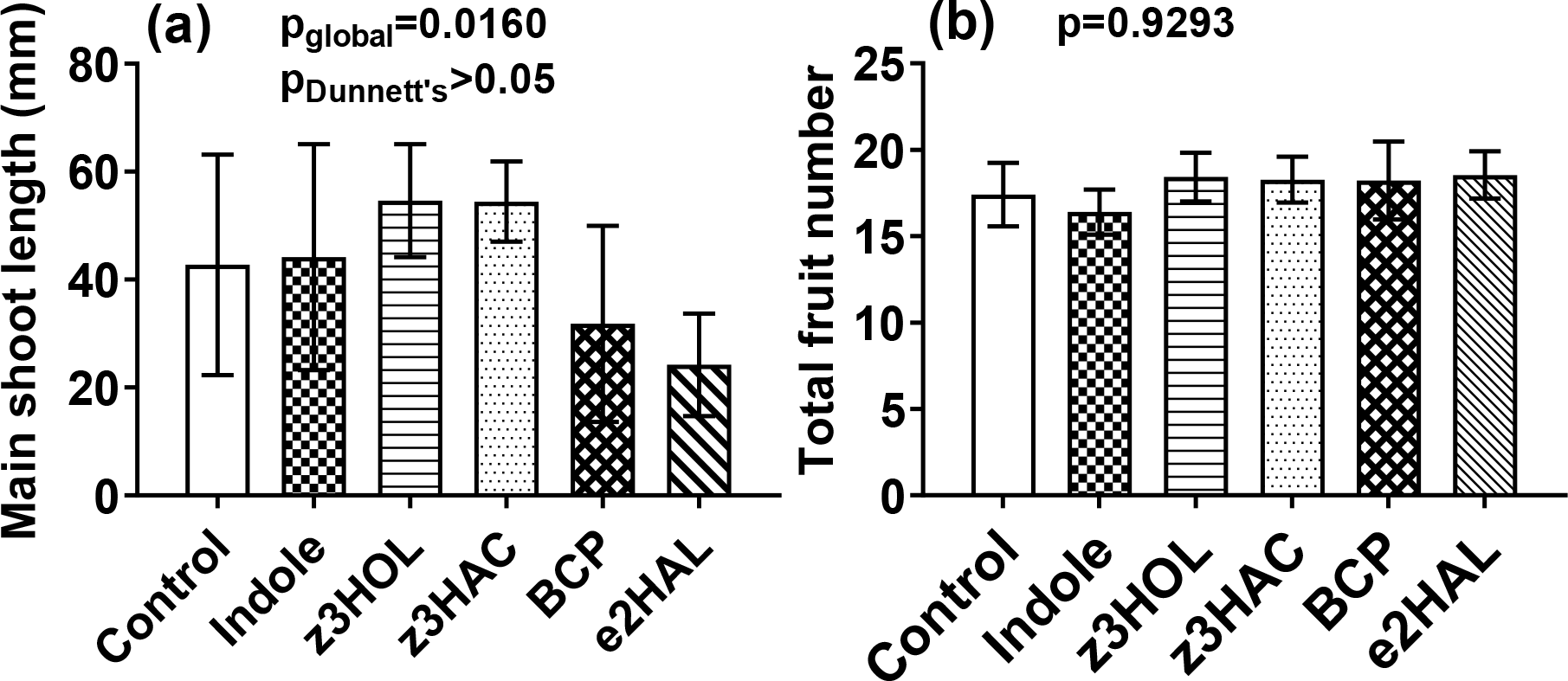
Seed exposure to plant derived volatiles have no effect on (a) Main shoot length and (b) Total fruit number of *M. truncatula* plants. Values are shown as means ± 95% CI (n=5-10) significance was calculated by one-way ANOVA followed by Dunnett’s post-hoc test.

**Fig. S3.**
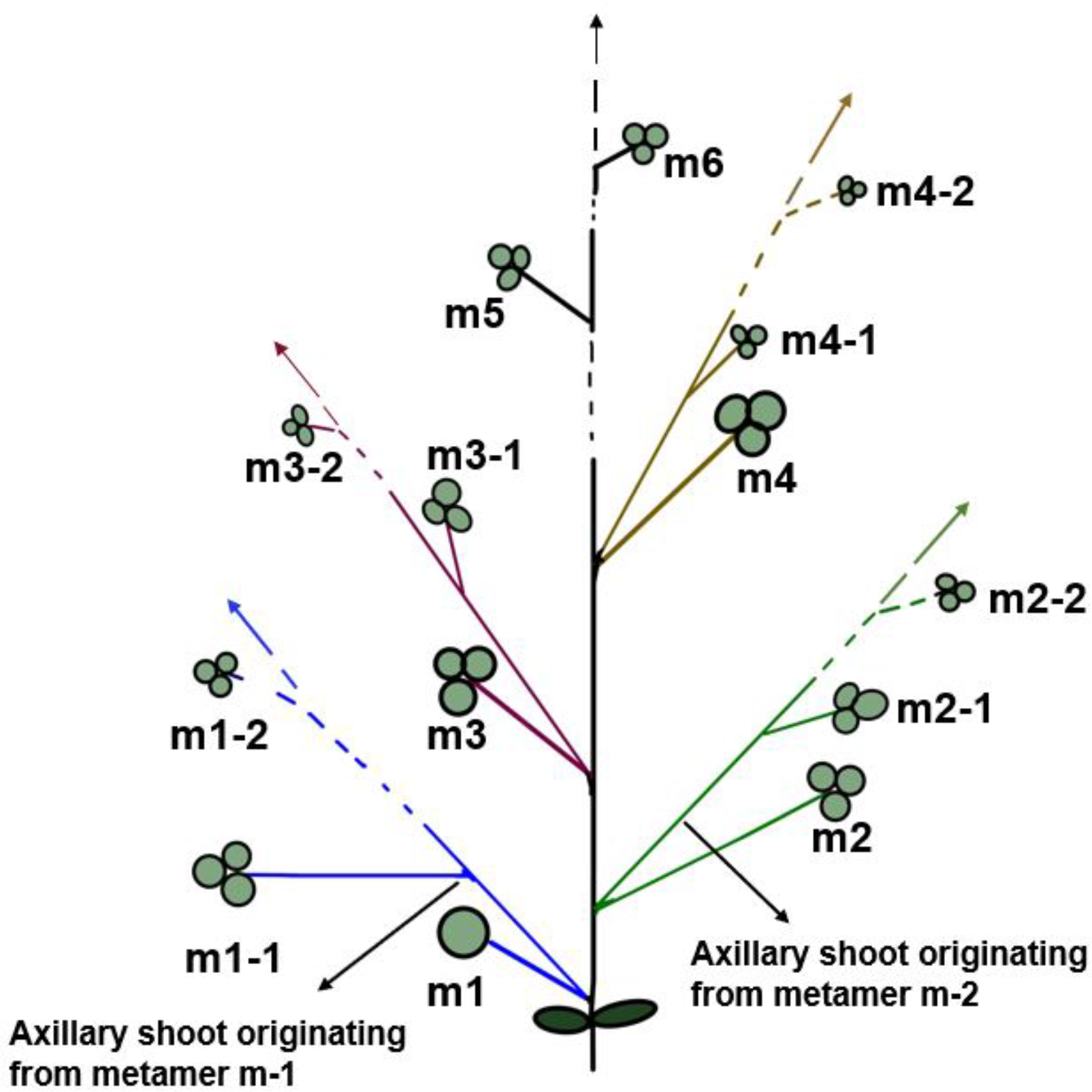
Pictorial representation of the numerical nomenclature coding system for vegetative growth of *M. truncatula*. Nomenclature coding started with unifoliate leaf as first metamers and subsequent trifoliate are labeled along the main shoot in ascending order. Axillary shoots are named as per the metamer of origin.

